# Hydraulic traits are more diverse in flowers than in leaves

**DOI:** 10.1101/461244

**Authors:** Adam B. Roddy, Guofeng Jiang, Kunfang Cao, Kevin A. Simonin, Craig R. Brodersen

## Introduction

The appearance of modern flowers was a turning point in the evolution of flowering plants, enhancing the intimacy and extent of interactions with their animal pollinators and increasing the likelihood of successful outcrossing (Crane *et al*., 1995; Fenster *et al*., 2004). Flowers evolved new morphological traits and rapidly diversified, effectively increasing the number of axes along which species could differentiate (Stebbins, 1951; Crepet & Niklas, 2009). Flowers are often noted for their incredible diversity of form, with high amounts of variation existing across phylogenetic scales, from within species to among major clades (Chartier *et al*., 2014, 2017; O’Meara *et al*., 2016), and these seemingly complex shapes can be formed through simple processes (Liang & Mahadevan, 2011).

Despite this obvious diversity in floral form, flowers all experience similar biophysical constraints of supplying resources during their development and maintaining a biomechanically functional structure on display for attracting pollinators and protecting the developing embryos. While the incredible diversity of floral morphologies and the ease with which certain complex shapes can be formed may imply that any form is possible, flowers are nonetheless constrained by their development and physiology (Berg, 1960; Strauss & Whittall, 2006; Roddy *et al*., 2013). Furthermore, flowers are produced and function in the context of the entire plant, and investment in flowers can often come at the cost of the function of vegetative organs (Galen, 1999; Galen *et al*., 1999; Lambrecht & Dawson, 2007; Lambrecht, 2013). The allocation of resources to vegetative growth or reproduction are critical components of plant life history strategy (Bazzaz *et al*., 1987). Yet the costs of reproduction are typically quantified solely as the biomass costs (Reekie & Bazzaz, 1987a, b), even though flower water costs can be high and can directly affect both short-term and long-term physiological function of leaves (Galen, 1999; Galen *et al*., 1999; Lambrecht & Dawson, 2007; Roddy & Dawson, 2012).

Despite the importance of flowers to both angiosperm ecology and evolution, even basic information about their physiological function is lacking (Gleason, 2018). Because flowers are predominantly heterotrophic and do not assimilate substantial amounts of carbon (but see: Galen *et al*., 1993), it is thought that they may not need to transpire large amounts of water, and the total amount of water transpired by flowers may be strongly linked to the environmental conditions during flowering (Blanke & Lovatt, 1993; Liu *et al*., 2017; Roddy *et al*., 2018). Indeed, among extant flowers, monocots and eudicots have much lower hydraulic conductance than early-divergent magnoliid flowers, suggesting that flowers have evolved towards lower hydraulic efficiency due possibly to a reduction of veins and stomata and to a decoupling of the developmental programs controlling hydraulic traits in leaves and flowers (Lipayeva, 1989; Roddy *et al*., 2013, 2016; Zhang *et al*., 2017). Yet these few traits do not entirely define the range of possible physiological strategies flowers may employ. For example, in lieu of efficiently transporting water from the stem to meet transpirational demands (i.e. relying on high *Kflower*), flowers may rely on high hydraulic capacitance to minimize water potential declines that may otherwise lead to embolism formation and spread (Chapotin *et al*., 2003; Zhang & Brodribb, 2017; Roddy *et al*., 2018), a strategy important to vegetative structures as well (Meinzer *et al*., 2003, 2009; McCulloh *et al*., 2014). Yet, there has not been an assessment of the diversity of hydraulic strategies of flowers. Furthermore, given the incredible morphological complexity and diversity of flowers–each flower is composed of multiple organs, and the developmental bases for these organs varies among species (Irish, 2009, 2017; Specht & Bartlett, 2009)–and that the physiological constraints of efficiently transporting high fluxes of water may have been relaxed in flowers (Roddy *et al*., 2016) flower hydraulic traits may not be constrained to varying along the same axes as leaves.

Here we measured pressure-volume relations of leaves and flowers of 22 species that include magnoliids, monocots, and eudicots from temperate and subtropical environments to quantify the variation in floral drought responses. Based on differences in lifespan and function, we predicted that flowers and leaves would differ in a few major ways. First, we predicted that flowers would have higher (less negative) turgor loss points (Ψ_tlp_) and higher hydraulic capacitance than leaves, reflecting a strategy of using hydraulic capacitance to minimize water potential declines (Chapotin *et al*., 2003; Roddy *et al*., 2018). Second, we predicted that flowers would not be as constrained in their bivariate scaling relationships as leaves because flowers need not maintain high hydraulic conductance. Third, we predicted that the convergence of traits associated with low hydraulic conductance (*Kflower*, vein and stomatal densities) among even widely divergent species (Roddy *et al*., 2013, 2016; Zhang *et al*., 2018) would allow other hydraulic traits associated with drought tolerance to vary more widely.

## Methods

### Plant material

Species were chosen to include a broad phylogenetic sampling and were selected based on the amenability of measuring water potentials on their flowers or inflorescences. Because the pressure chamber requires that a minimum length of the pedicel extend through the enclosing gasket, many species could not be measured. While we could have included short segments of subtending shoots, we avoided this because the inclusion of stems could have introduced artifacts by biasing water content measurements. Plants were grown outdoors, under well-watered conditions, in botanical gardens and on university campuses (Table 1). These sites and species included both temperate (Marsh Botanical Garden, New Haven, CT, USA; Arnold Arboretum, Jamaica Plain, MA, USA; University of California Botanic Garden, Berkeley, CA, USA; campus of Yale University) and subtropical (campus of Guangxi University, Nanning, China) sites. Flowering shoots were collected from at least three individuals per species and immediately recut underwater in the early morning and allowed to rehydrate for at least 30 minutes before individual flowers or leaves were excised for measurement. Potted plants of *Rosa sp.*, *Anthurium andraeanum*, and *Dendrobium sp.* were maintained well-watered prior to sampling. The morphology of monocot leaves precluded measurements in the pressure bomb, with the exception of *Anthurium andraeanum*. For this species the entire inflorescence, including both the spathe and the spadix were measured.

**Table 1.** List of species and their collection locations.

| Species | Family | habit | Collection location |
| --- | --- | --- | --- |
| magnoliids |  |  |  |
| *Calycanthus occidentalis* | Calycanthaceae | shrub | UC Botanical Garden |
| *Calycanthus floridus* | Calycanthaceae | shrub | Marsh Botanical Garden |
| *Calycanthus chinensis* | Calycanthaceae | shrub | UC Botanical Garden |
| *Liriodendron tulipifera* | Magnoliaceae | tree | Marsh Botanical Garden |
| *Magnolia sieboldii* | Magnoliaceae | tree | Arnold Arboretum |
| *Magnolia stellata* | Magnoliaceae | tree | Marsh Botanical Garden |
| *Magnolia loebneri* | Magnoliaceae | tree | Marsh Botanical Garden |
| monocots |  |  |  |
| *Anthurium andraeanum* | Araceae | shrub | Guangxi University |
| *Lilium lancifolium* | Liliaceae |  | Arnold Arboretum |
| *Dendrobium sp.* | Orchidaceae |  | Guangxi University |
| *Hemerocallis sp.* | Xanthorrhoeaceae |  | Marsh Botanical Garden |
| eudicots |  |  |  |
| *Clematis sp.* | Ranunculaceae | liana | Marsh Botanical Garden |
| *Aquilegia sp.* | Ranunculaceae | shrub | Marsh Botanical Garden |
| *Ceiba speciosa* | Malvaceae | tree | Guangxi University |
| *Rosa sp.* | Rosaceae | shrub | Guangxi University |
| *Bauhinia blakeana* | Fabaceae | tree | Guangxi University |
| *Calliandra*<br>*haematocephala* | Fabaceae | shrub | Guangxi University |
| *Bougainvillea glabra* | Nyctaginaceae | shrub | Guangxi University |
| *Cornus florida* | Cornaceae | tree | Marsh Botanical Garden |
| *Bidens pilosa* var.<br>*radiata* | Asteraceae | herb | Guangxi University |
| *Stewartia*<br>*pseudocamellia* | Theaceae | tree | Arnold Arboretum |
| *Rhododendron sp.* | Ericaceae | shrub | Marsh Botanical Garden |

### Measurement of pressure-volume parameters

Shoots were allowed to rehydrate and water potentials equilibrate for at least two hours before individual flowers or leaves were excised and initial water potentials measured. Initial water potentials were always higher than -0.15 MPa. Following standard methods, pressure-volume curves were constructed for each sample by repeatedly measuring the bulk water potential using a Scholander-style pressure chamber (0.01 MPa resolution; PMS Instruments, Albanay, OR, USA) and subsequently measuring the fresh mass to determine the relationship between water potential and water content (Sack *et al*., 2010, 2011). Because rates of water potential change are nonlinear and water potential initially declines rapidly, specimens were briefly exposed to ambient laboratory air and then enclosed in humidified plastic bags to allow equilibration of water potentials among tissue types. After the point of turgor loss, the duration of exposure to a dry laboratory atmosphere was lengthened to allow sufficient declines in water potential. The pressure chamber was kept humidified with wet paper towels to prevent evaporation during the water potential measurement. The balancing pressure was determined by slowly increasing the pressure inside the chamber until water was expressed at the cut petiole or pedicel surface, at which time the pressure inside the chamber was slowly relaxed to ambient pressure. Immediately afterward, the specimen was weighed. After the conclusion of the measurements, each specimen was oven-dried at 70°C for at least 72 hours prior to determining dry mass. Contrary to prior measurements on leaves, we expressed pressure-volume parameters on dry mass basis, rather than on a surface area basis, to facilitate comparisons between flowers and leaves, because the complex morphologies of flowers and their high degree of shrinkage prevented accurate measurements of surface area after pressure-volume measurements were complete.

### Phylogeny

We used Phylomatic (v 3.0) to generate a family-level supertree using the R package ‘brranching’. This supertree is in good agreement with the most recent understanding of the relationships between angiosperm families (Group, 2016). Nodes in the tree were dated using age estimates from (Magallón *et al*., 2015), and all branch lengths smoothed using the function ‘bladj’ in Phylocom (Webb *et al*., 2008), following methods of previous studies (Roddy *et al*., 2013; Simonin & Roddy, 2018). This dated phylogeny was used in all subsequent phylogenetic analyses. For comparisons of trait values between leaves and flowers (phylogenetic paired t-tests), data were not available for monocot leaves, and so the phylogeny was pruned of these species for these analyses.

### Data analysis

All statistical analyses were performed in R (v. 3.5.0). Two metrics of phylogenetic signal were calculated for each trait, Pagel’s λ and Abouheif’s *C_mean_* because of the robustness of these two measures (Münkemüller *et al*., 2012), using the package ‘phylosignal’ (Keck *et al*., 2016). Phylogenetic paired t-tests (Revell, 2012) were used to compare differences in each trait between leaves and flowers. Because the leaves of three monocot species were not measured, these species were entirely omitted from paired t-tests, although values for their traits are reported in Figure 1.

**Figure 1.**
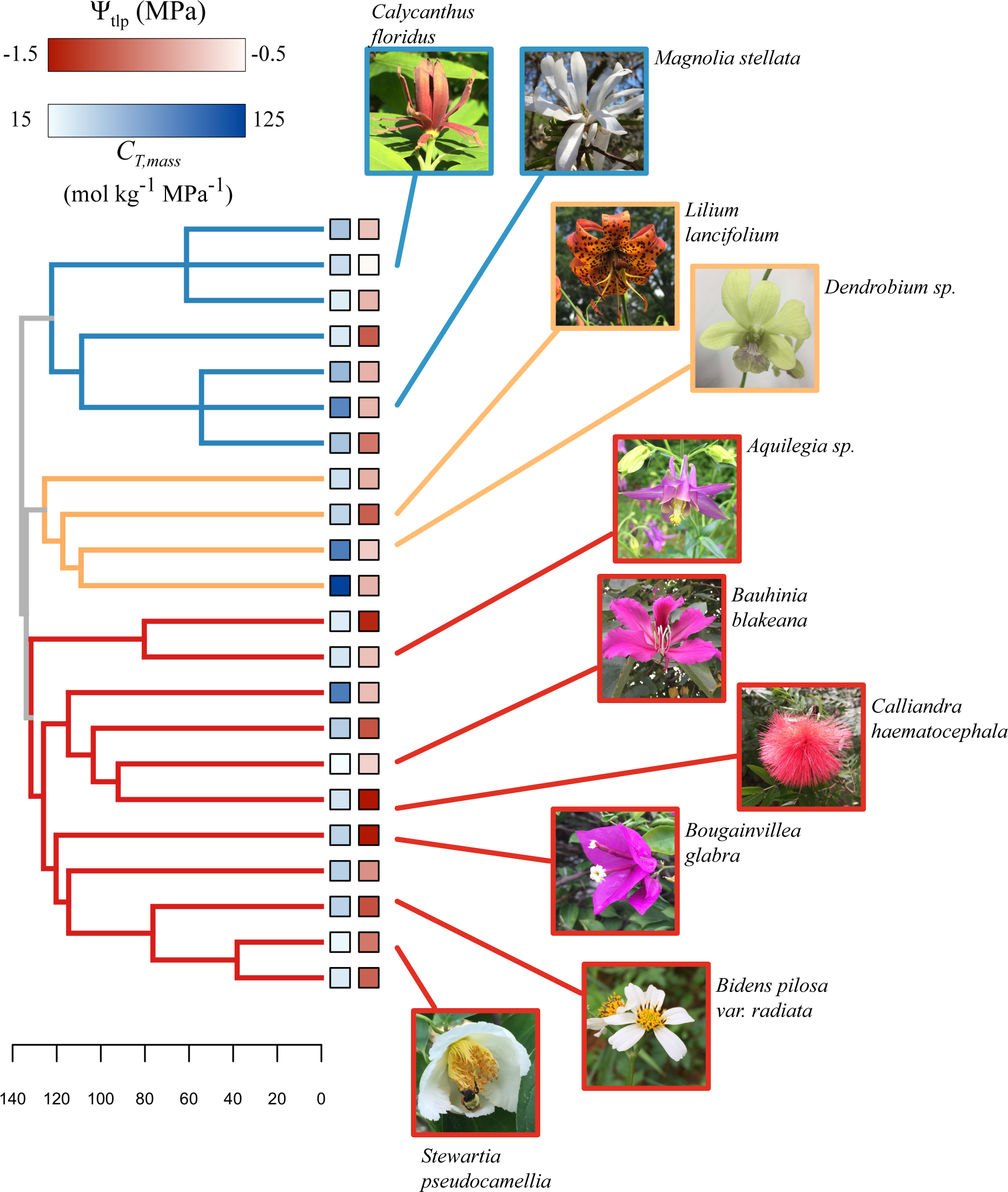
Morphological diversity of the species sampled. Phylogenetic relationships of the species sampled with values of hydraulic capacitance before turgor loss (*C_T, mass_*) and the water potential at turgor loss (Ψ_tlp_) for flowers mapped on the tips. Photos of species sampled highlight the morphological diversity. Branches are colored according to clade (blue = magnoliids, orange = monocots, red = eudicots), and these colors are used in all subsequent figures. All photos taken by ABR.

Standard major axis regression was used to determine scaling relationships between traits (the function ‘sma’ in the package *smatr*; Warton *et al*., 2012)). We tested for differences in scaling slopes, intercepts, and whether there were shifts along scaling axes and report relevant statistics for each of these tests: for comparisons of slopes the likelihod ratio test (LRT) statistic is used but for differences in elevation and for shifts along common slopes the Wald statistic is used. For all comparisions except for the relationship between Ψ_sft_ and Ψtlp, data were log-transformed. However, for visualization purposes data are plotted in arithmetic space with regression lines appropriately transformed. In figure insets, data are plotted in log space for comparison.

Principal components were calculated using the function ‘prcomp’ on the data for each individual specimen measured to determine the loadings of traits. Means of the PC scores for each species and structure were calculated to compare the total multivariate space occupied by flowers and leaves.

## Results

### Traitwise differences between leaves and flowers

Although the range of each trait overlapped for flowers and leaves, paired t-tests correcting for shared evolutionary history revealed that flowers and leaves differed significantly in almost every trait (Figure 2). Flowers had significantly higher *SWC* (t = 6.18, P < 0.0001), *C_T, mass_* (t = 3.96, P < 0.01), *C_t, mass_* (t = 2.86, P = 0.01), *N_s, mass_* (t = 3.46, P < 0.01), Ψ_tlp_ (t = 4.30, P < 0.001), and Ψ_sft_ (t = 4.76, P < 0.001) but lower ∊_bulk_ (t = 2.30, P = 0.04). There were no significant differences between structures in *RWC_tlp_*.

**Figure 2.**
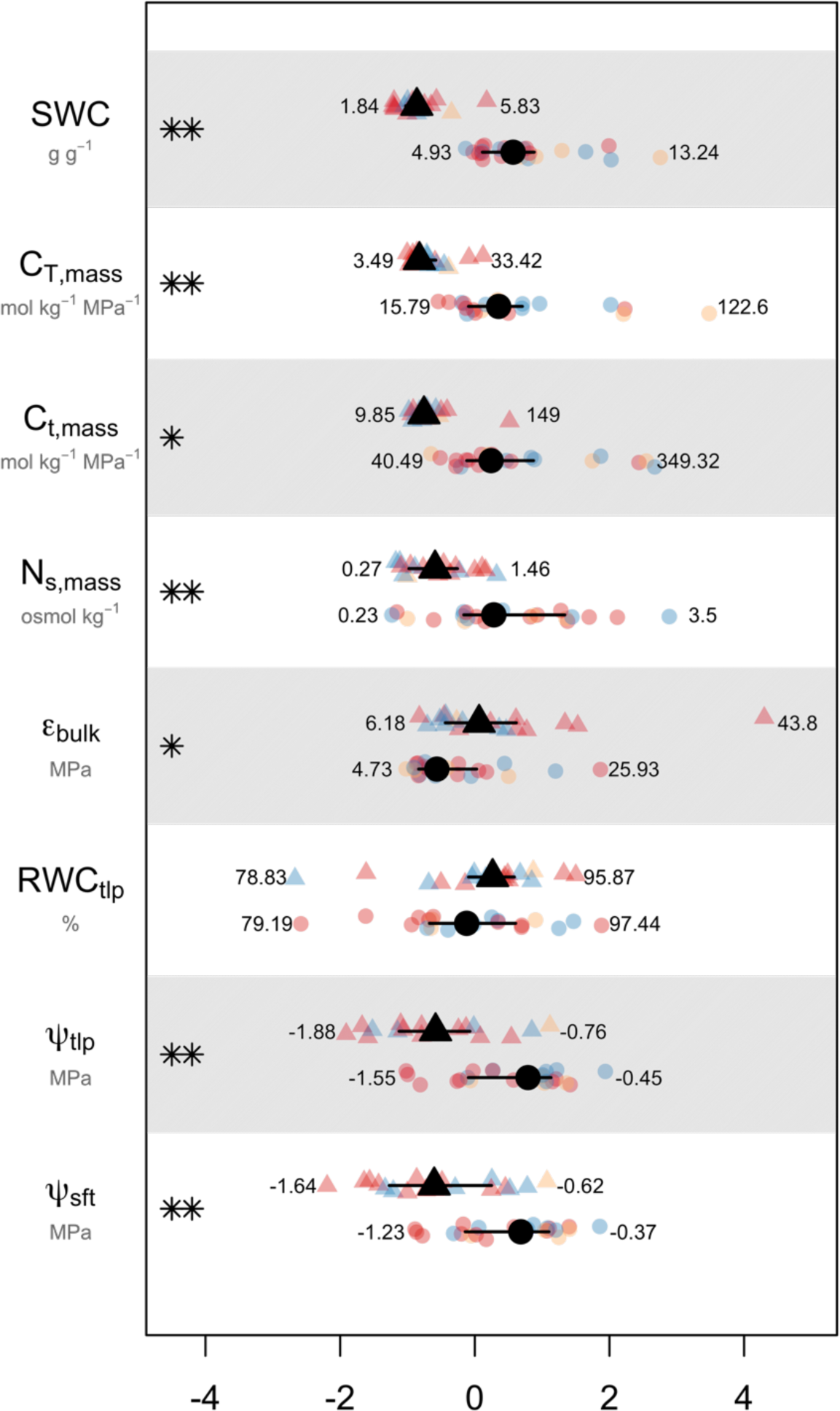
Standardized differences between leaves (triangles) and flowers (circles) in pressure-volume traits. Standardized differences between leaves (triangles) and flowers (circles) in hydraulic traits. Black points and lines indicate medians ± interquartile ranges for each structure. Colored points are mean values for each species, colored by clade (blue = magnoliids, orange = monocots, red = eudicots). Numbers indicate the maximum and minimum species means of each trait for each structure. Asterisks indicate significant differences between structures in phylogenetically-controlled t-tests: * α = 0.05; ** α = 0.05

### Coordination between traits

A single slope described the relationship between *CT, mass* and *CT, mass* across species and structures (R^2^ = 0.78, P < 0.0001; Figure 3a), and flowers were shifted along this common slope towards higher capacitance values (Wald statistic = 53.84, P < 0.0001). Capacitance both before (*C_T, mass_*) and after (*C_T, mass_*) turgor loss was strongly predicted by *SWC* (Figure 3b, c). The relationship between *SWC* and *C_T, mass_* was described by a common slope and intercept among leaves and flowers (R^2^ = 0.81, P < 0.0001; slope: LRT = 0.16, P = 0.69; intercept: Wald statistic = 3.46, P = 0.06), although flowers were shifted along this common line (Wald statistic = 66.46, P < 0.0001). Similarly, *SWC* predicted *C_T, mass_* across species and structures with a single slope (R^2^ = 0.76, P < 0.0001; slope: LRT = 0.03, P = 0.87; intercept: Wald statistic = 3.47, P = 0.06), although flowers were shifted along this common axis (Wald statistic = 63.58, P < 0.0001).

**Figure 3.**
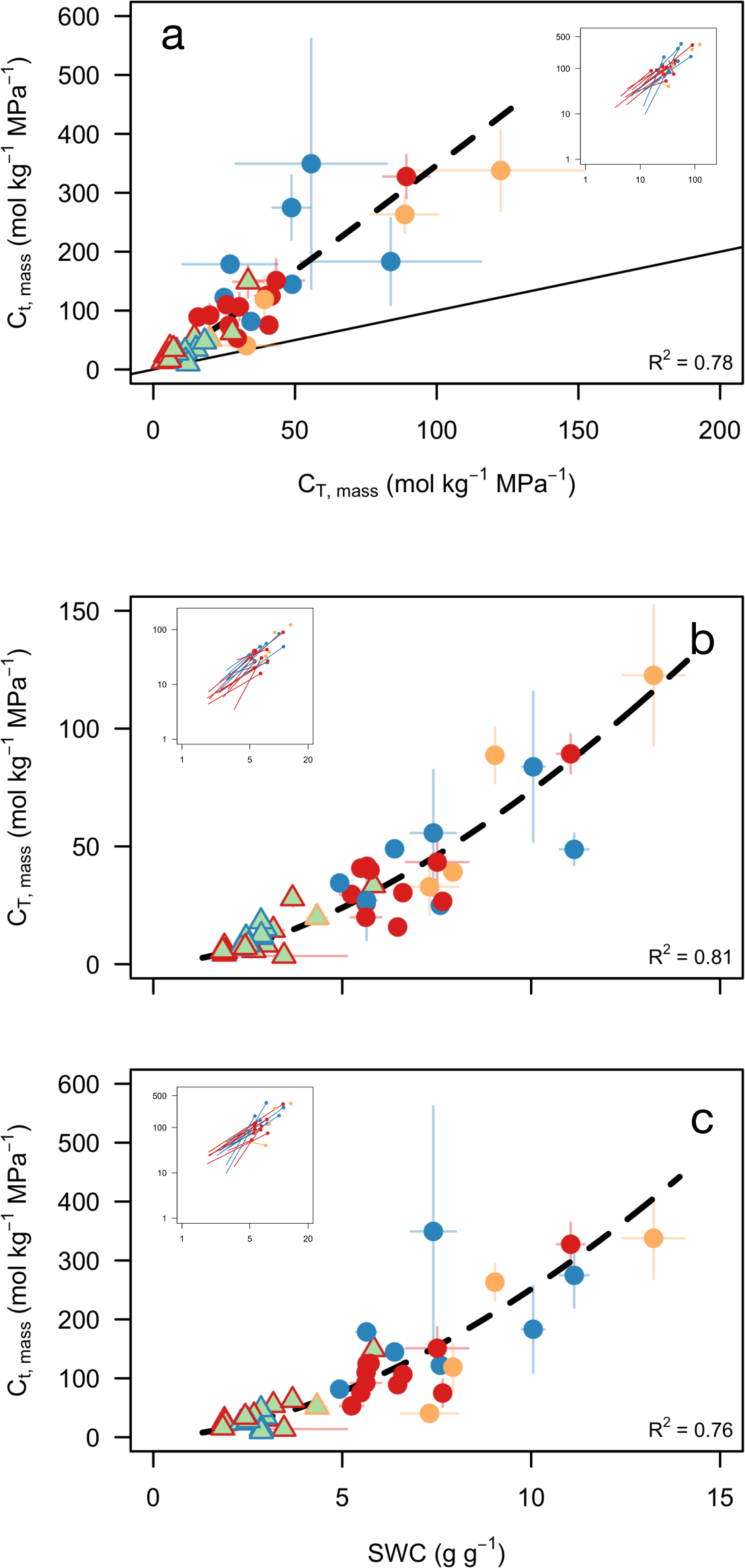
Relationships between saturated water content (SWC) and hydraulic capacitance before (CT, mass) and after (Ct, mass) the turgor loss point. Relationships between saturated water content (*SWC*), hydraulic capacitance before turgor loss (*C_T, mass_*), and hydraulic capacitance after turgor loss (*C_t, mass_*). Insets show log-log relationships and lines connect conspecific leaves and flowers. Dashed black lines are standard major axis regressions of log-transformed data. (a) Hydraulic capacitance after turgor loss is higher than before turgor loss. The solid line is the 1:1 line, and the dashed line is the standard major axis regression. (b, c) *SWC* is a strong predictor of hydraulic capacitance both before and after turgor loss.

There was a highly significant, negative relationship between *CT, mass* and the bulk modulus of elasticity (Figure 4a), with slope and elevation tests revealing that leaves and flowers having statistically indistinguishable slopes (leaves: -1.30, R^2^ = 0.77, P < 0.0001; flowers: -1.10, R^2^ = 0.59, P < 0.001; slope test: LRT = 0.79, P = 0.37) but different intercepts (Wald statistic = 69.63, P < 0.0001). Furthermore, flowers are shifted along this common scaling axis (Wald statistic = 26.83, P < 0.0001).

**Figure 4.**
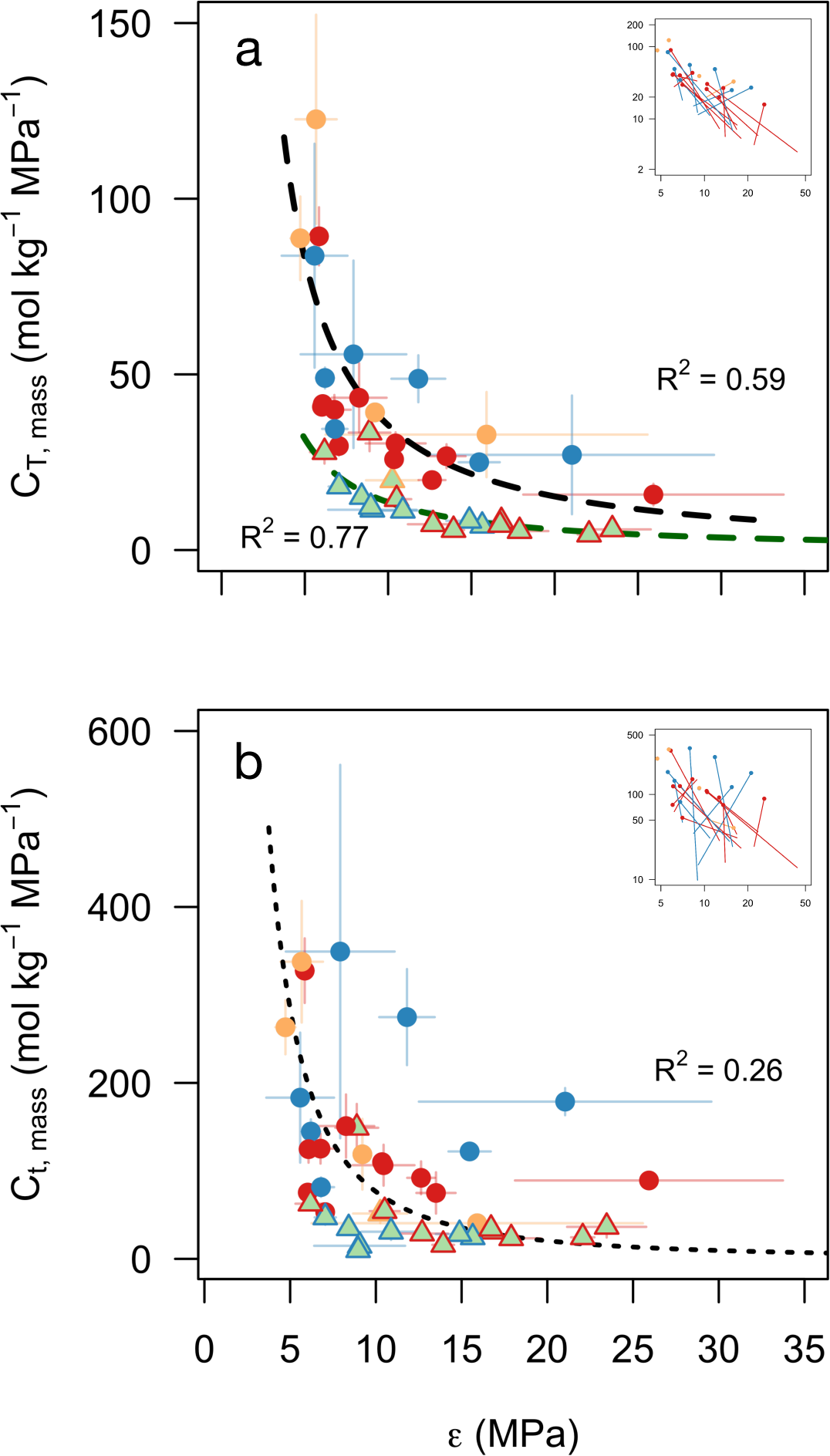
Relationships between the bulk modulus of elasticity and hydraulic capacitance before (CT, mass) and after (Ct, mass) the turgor loss point. Relationships between the bulk modulus of elasticity and hydraulic capacitance before turgor loss (*C_T, mass_*), and hydraulic capacitance after turgor loss (*C_T, mass_*). Insets show log-log relationships, and lines connect conspecific leaves and flowers. Dashed black lines in (a) indicate standard major axis regressions of log-transformed data for leaves and for flowers separately. Dotted line in (b) is fit through all the data.

The relationship between elasticity and *C_T, mass_* showed a similar, significant, negative relationship (Figure 4b). However, there was no significant difference in slopes between structures (LRT = 0.009; P = 0.92), and a single slope existed among structures (R^2^ = 0.26, P < 0.001).

There was no significant difference between structures in the relationship between Ψ_sft_ and Ψtlp (R^2^ = 0.94, P < 0.0001; Figure 5). While there was no significant difference between structures in intercepts (Wald statistic = 0.08, P = 0.78), flowers were shifted towards higher values in both traits (Wald statistic = 15.10, P < 0.001), although phylogenetically controlled t-tests revealed no significant difference between leaves and flowers in either of these traits (Figure 2).

**Figure 5.**
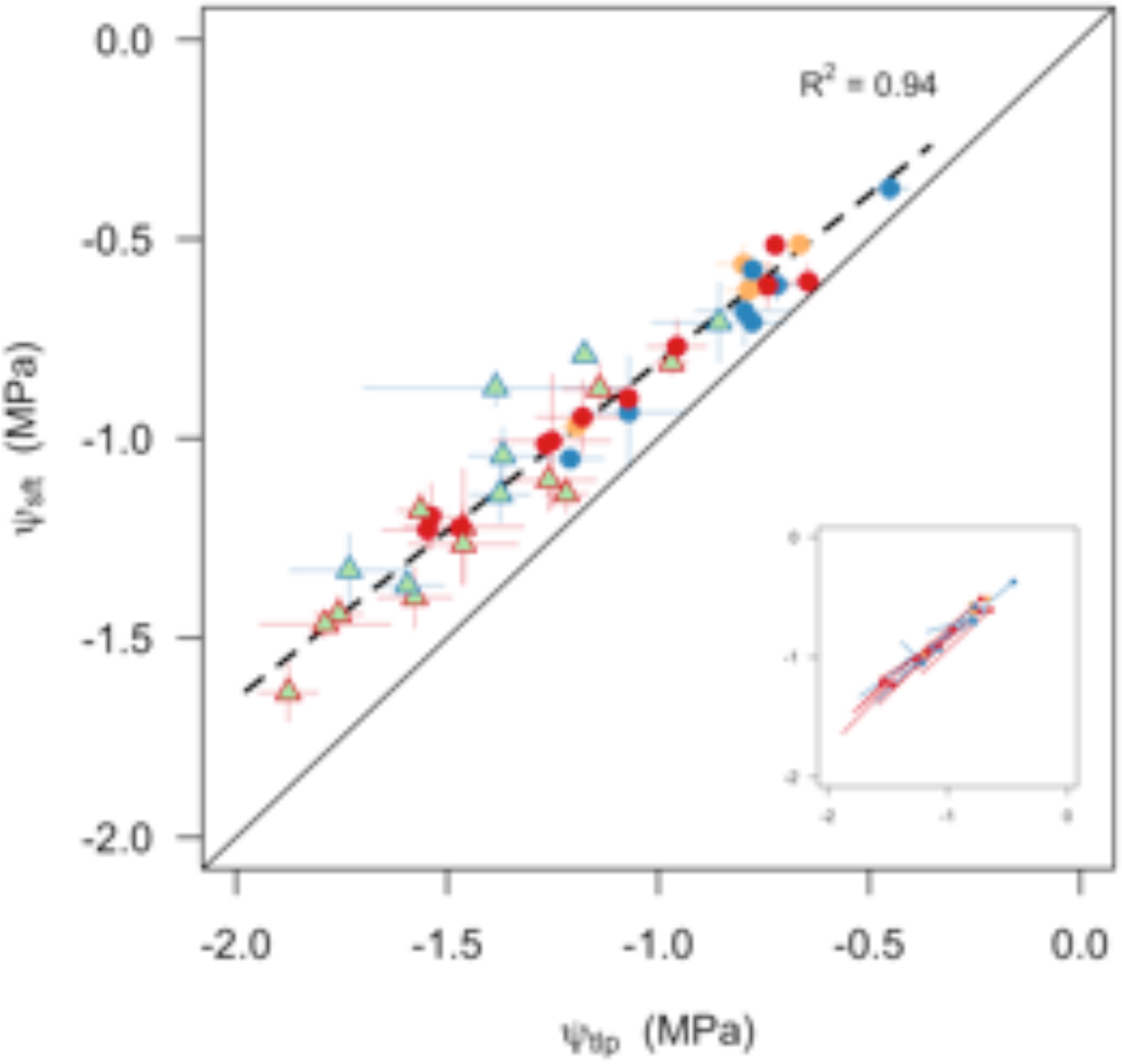
Relationship between the osmotic potential at full turgor (Ψ_sft_) and the water potential at the point of turgor loss (Ψ_tlp_). The relationship between Ψ_sft_ and Ψ_tlp_ is the same for leaves and flowers. The dashed black line represents the standard major axis regression for both leaves and flowers. The solid black line is the 1:1 line. Lines in the inset connect conspecific leaves and flowers.

### Multivariate analysis of traits

In multivariate space, first to principal component axes explained 51% and 27% of the variation among all samples (Figure 6a, b). Differences in the first axis (PC1) were driven by a tradeoff between ∊_bulk_ and traits related to water content and discharge (*SWC*, *C_T, mass_*, *C_T, mass_*). Flowers and leaves differed in the regions of trait space they occupied, consistent with pairwise differences in traits (Figure 2). There was little overlap between flowers and leaves in the first two PC axes, and flowers occupied a larger volume of trait space than did leaves (Figure 6b).

**Figure 6.**
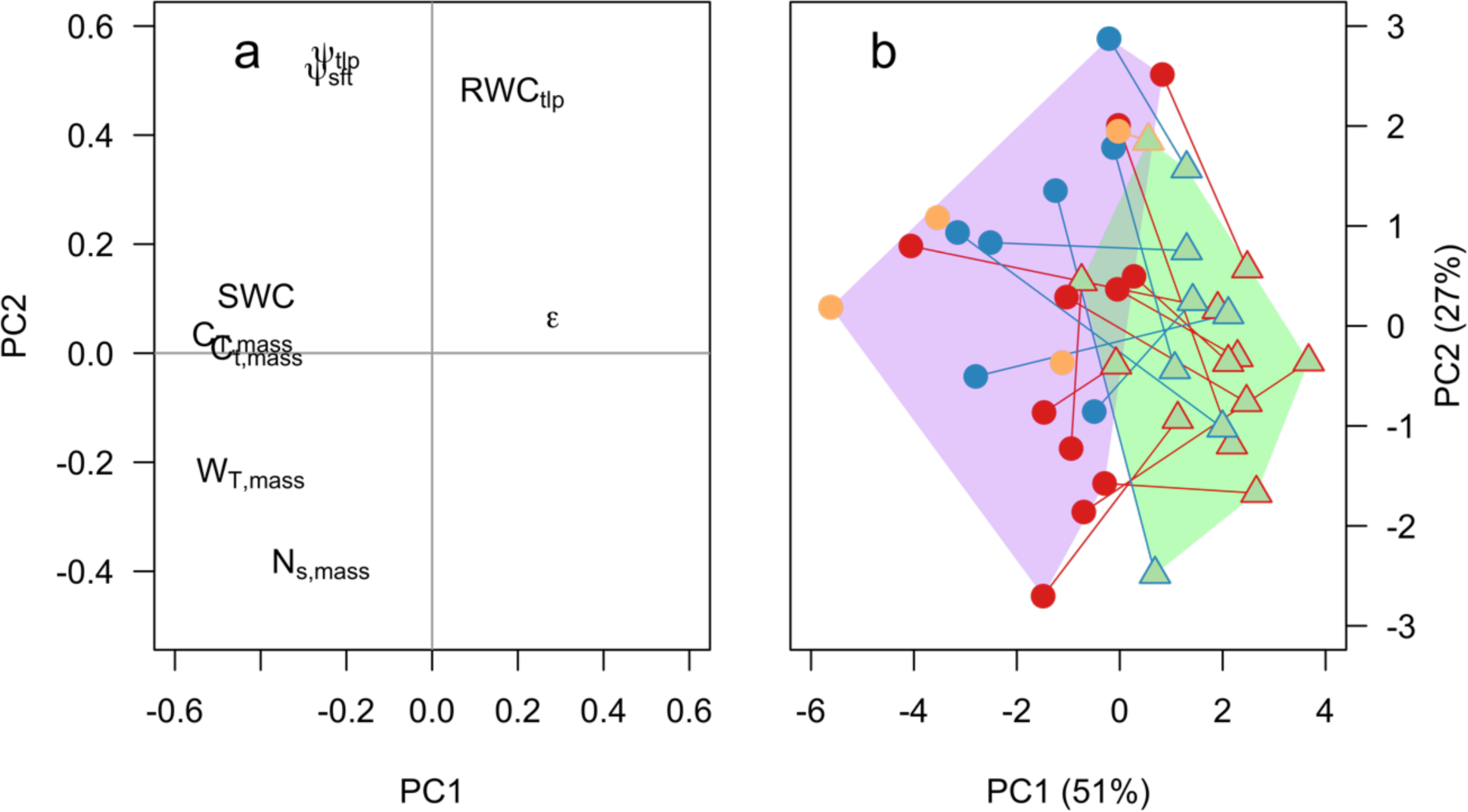
(a) Loadings of each trait in the first two principal component axes. (b) Principal component scores for leaves and flowers. Results of principal components analysis performed on raw data for both leaves and flowers. (a) Loadings of the first two PC axes explain a total of 79% of the variation in leaf and flower pressure-volume parameters. (b) Mean scores for species and structures in the first two PC axes. Lines connect conspecific leaves and flowers, and colors indicate phylogenetic clade. The shaded regions indicate the total volume of trait space occupied by leaves (green) and flowers (purple).

## Discussion

The diversity and evolution of flowers has long intrigued scientists and naturalists alike, and early work by Kohlreuter and Sprengel pointed to the fundamental role of flowers in plant reproduction (Sprengel, 1793, 1996; Vogel, 1996). More recent studies have characterized the strength of these coevolutionary relationships with animal pollinators and how they have driven floral form (Fenster *et al*., 2004; Whittall & Hodges, 2007). But flowers also suffer antagonistic relationships with herbivores (Strauss, 1997) and are constrained by environmental and physiological factors associated with their production and maintenance (Bazzaz *et al*., 1987; Reekie & Bazzaz, 1987a, b, c; Galen, 1999), and the existence of diverse, often opposing agents of selection may help to promote variation within species (Strauss & Whittall, 2006), as well as providing more numerous axes along which species can differentiate. How these non-pollinator agents of selection, such as physiological traits linked to water supply and turgor maintenance, vary among species and may have contributed to floral evolution and diversification has remained largely unstudied. Yet, recent studies have shown strong phylogenetic trends in floral hydraulic traits (Roddy *et al*., 2016), suggesting that floral diversification may be linked to innovations in floral physiology, just as innovations in leaf anatomy and physiology among the angiosperms are associated with changes in diversity and dominance (Simonin & Roddy, 2018).

Here we show that while similar scaling relationships govern floral and foliar hydraulic traits, flowers encompass a wider diversity of drought strategies than leaves. Despite being comparatively ephemeral, the hydraulic constraints that govern leaf hydraulic architecture and physiological function may be relaxed in flowers. While leaves were evolving towards increasing the densities of veins and stomata with the effect of increasing hydraulic conductance and gas exchange rates (Boyce *et al*., 2009; Brodribb & Feild, 2010; Feild *et al*., 2011; Boer *et al*., 2012; Simonin & Roddy, 2018), flowers seem to have been evolving towards reducing vein and stomatal densities with the effect of reducing hydraulic conductance (Roddy *et al*., 2013, 2016; Zhang *et al*., 2018). Such developmental modularity of even the same traits (Roddy *et al*., 2013) was likely an important innovation enabling the independent diversification and specialization of each of these complex structures.

### Similar scaling relationships govern leaf and flower hydraulic architecture

Consistent with our first prediction flowers had significantly higher hydraulic capacitance both before and after turgor loss than leaves, as well as higher turgor loss points than leaves (Figure 2). The relationship between Ψ_tlp_ and Ψ_sft_ was the same for leaves and for flowers, suggesting that methods for rapidly assessing turgor loss points in leaves (Bartlett *et al*., 2012) may be applicable to flowers as well. In addition to hydraulic capacitance, the traits showing the largest differences between flowers and leaves were *SWC* and *Ns, mass*, with flowers having higher trait values for all of these traits. Indeed, *SWC* was a strong predictor of hydraulic capacitance before and after turgor loss (Figure 3), which together with the ∊_bulk_ provided the major axis of differentiation between structures (Figure 6).

Consistent with our second hypothesis, hydraulic traits were generally more variable among flowers than among leaves, although flowers and leaves obeyed similar scaling relationships between traits (Figures 3-5). For example, the differences in hydraulic capacitance were driven by consistently higher *SWC* in flowers compared to conspecific leaves (Figure 3b, c insets). However, in other cases, while scaling slopes were equivalent for leaves and flowers, the intercepts differed; for a given ∊_bulk_, flowers had higher *C_T, mass_*. This difference in intercepts between structures is due to differences in *SWC* (Figure 3). But where is this extra water per unit dry mass stored in flowers? First, cells in flowers could be larger such that the ratio of vacuole volume to cell wall is higher, and some evidence suggests that epidermal pavement cells and guard cells may be larger in petals than in leaves (Zhang *et al*., 2018). Additionally, the higher *SWC* of flowers could be due to the presence of extracellular water stored in the form of mucilage, which has been reported previously in flowers (Chapotin *et al*., 2003) and which we frequently observed in various floral structures (e.g. petals, gynoecia) upon excision and dissection. Furthermore, the presence of extracellular mucilage has been linked to increased hydraulic capacitance in both leaves (Morse, 1990) and flowers (Chapotin *et al*., 2003), suggesting that storing water as mucilage may be an effective way of avoiding water potential declines. These results provide strong support that the basic rules governing the hydraulic architecture of flowers are the same as or similar to those governing leaf hydraulic architecture but that leaves are clustered towards the extreme ends of these trait spectra.

### A critical role for hydraulic capacitance

Although the relationships between traits are similar for flowers and for leaves, flowers nonetheless diverge in their positions along these axes towards having higher water contents and hydraulic capacitance. Hydraulic capacitance–the amount of water discharged for a given change in water potential–may be a key functional trait in flowers that are otherwise incapable of efficiently moving water to meet their transpirational demands. High hydraulic capacitance can serve three major functions: (1) it can supply water for transpiration without the need for continuous water import, (2) it minimizes water potential declines that may otherwise lead to turgor loss and wilting, and (3) it can help delay embolism formation and spread. All of these functions may explain the universally high hydraulic capacitances found in flowers.

Hydraulic capacitance has long been recognized as a functionally important trait in succulent plants that is linked to overall ecological strategy as well as the structure and function of specific organs (Nobel & Jordan, 1983; Arakaki *et al*., 2011; Ogburn & Edwards, 2013), but in non-succulent plants, hydraulic capacitance is only more recently being acknowledged as important to the maintenance of a functioning hydraulic system (Meinzer *et al*., 2003, 2009; McCulloh *et al*., 2014; Roddy *et al*., 2018). Among co-occurring desert species leaf hydraulic capacitance can vary more than three orders of magnitude, allowing species with succulent leaves to support their transpirational demands solely from stored water for two to three orders of magnitude longer than non-succulent species (Nobel & Jordan, 1983). The continuous discharge of water from storage components can decouple water uptake from water loss, effectively preventing steady-state transpiration (Hunt & Nobel, 1987), which is especially important for the measurement of gas exchange and isotope fluxes (Simonin *et al*., 2013). Buffering the effects of transpiration may be important in flowers, which have low vein densities and hydraulic conductance, and, therefore, may not be able to continuously supply enough water for transpiration (Roddy *et al*., 2016, 2018). Furthermore, the high hydraulic capacitance of reproductive organs has the potential to buffer water potential variation in the stem and leaves: diurnal declines in Ψstem can drive water flow from fruits back into the stem and be replaced nocturnally (Higuchi & Sakuratani, 2006).

Hydraulic capacitance (*C_T, mass_*) can also compensate for the high Ψ_tlp_ common among flowers. By allowing water content to decline with minimal effect on water potential, a high hydraulic capacitance can help delay water potential declines that lead to turgor loss (Morse, 1990; Meinzer *et al*., 2009; Roddy *et al*., 2018). With few stomata and limited control over them (Lipayeva, 1989; Roddy *et al*., 2016, 2018; Zhang *et al*., 2018), flowers could rapidly lose turgor without the conductive capacity to match their hydraulic supply to their water loss. Furthermore, in large, showy flowers, which must stay turgid to remain on display for animal pollinators but which have low investment in dry mass, avoiding turgor loss may be particularly important because visual wilting more directly accompanies turgor loss than it does in leaves, which have long-lived carbon-based support structures. Thus, capacitance may be critical in maintaining the biomechanical performance of flowers.

Finally, hydraulic capacitance can delay embolism formation and spread. Flowers and leaves alike had higher hydraulic capacitance after turgor loss (*C_T, mass_*) than before turgor loss (*C_T, mass_*; Figure 3a). After stomatal closure, aerial structures have few physiological mechanisms to prevent continued water loss and must rely instead on anatomical features. The epidermal conductance when stomata are fully closed (*g_min_*) has been proposed as an important trait conferring drought tolerance (Choat *et al*., 2018), and *C_T, mass_* may be another important trait that can prevent embolism formation and spread. Low density leaves with a low modulus of elasticity shrink substantially after turgor loss (Scoffoni *et al*., 2014), which reduces airspace porosity and brings hydrated cells into closer contact. In extreme cases, the tissue may shrink so much that no air remains in the tissue volume that could seed into the xylem and cause embolism spread. In this case, hydraulic capacitance after turgor loss may be due to continued discharge of intracellular water into the xylem. Whether this occurs in leaves is unclear, but some imaging results suggest that shrinkage may influence leaf vulerability to embolism (Scoffoni *et al*., 2017b, a). However, flowers have substantially lower tissue densities, and even *Calycanthus* tepals which have a high dry mass per area, revealed that almost all of the internal airspace disappeared after turgor loss, which may play a critical role in making these flowers so resistant to embolism in the xylem (Roddy *et al*., 2018). Maintaining xylem function after turgor loss would increase their capacity to rehydrate, potentially explaining why rehydration rates of flowers do not differ from those of leaves (Roddy and Brodersen, in prep.).

### Axes of floral physiological diversity

In contrast to previous results showing that there is strong phylogenetic signal in hydraulic traits of flowers (Roddy *et al*., 2016), the traits presented here lack similar phylogenetic structure and exemplify the diversity of extant flowers (Figure 6, Table 2). In fact, almost every trait was more variable among flowers than among leaves (Figure 2), which was reflected in the greater variation in multivariate trait space among flowers (Figure 6). However, despite this greater variation, flowers occupied a nearly distinct region of multivariate trait space, comapred to leaves. Leaves of only three species and flowers of only two species existed in the region of trait space common to leavs and flowers. The leaves and flowers of the monocot *Anthurium andraeanum*, which were very similar to each other, both existed in the overlapping region of trait space, and the other leaves and flowers in overlapping multivariate space were, surprisingly, all eudicots. Some have suggested that extant, basal angiosperm flowers, including those of the magnoliids, are more similar physiologically to leaves than are flowers of the eudicots (Feild *et al*., 2009a, b). However, the pressure-volume traits presented here show that the flowers most similar to leaves predominantly belong to the monocots and eudicots (Figure 6), even though basal angiosperm flowers have traits consistent with maintaining high transpiration rates (Roddy *et al*., 2016, 2018). This seeming paradox points to the importance of examining the full suite of traits associated with maintaining water balance because plant strategies are complex and rarely encapsulated by a single variable.

**Table 2.**
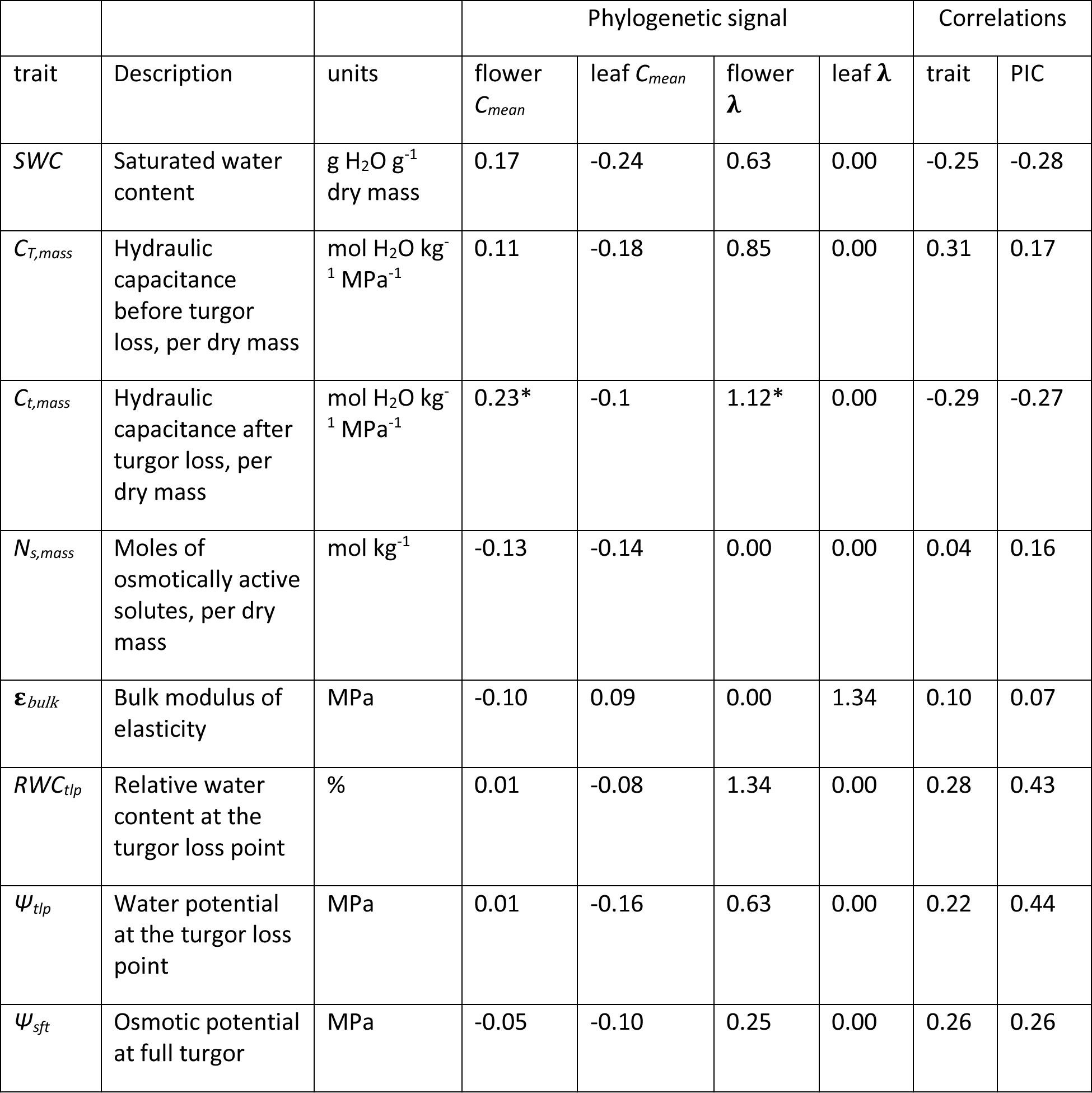
List of traits, units, their phylogenetic signal for leaves and for flowers, and the trait and phylogenetic independent contrast (PIC) correlations of each trait between leaves and flowers. *P < 0.05

While the interspecific variability in the floral hydraulic traits presented here is high, magnoliids are no less variable than monocots and eudicots (Figure 6). Detailed studies of the water relations of magnoliid flowers have shown that despite having both high hydraulic capacitance and hydraulic conductance that can exceed that of their conspecific leaves, these flowers are prone to wilting (Feild *et al*., 2009b; Roddy *et al*., 2018). Although reduced *K_flower_* among monocot and eudicot lineages is not necessarily associated with increased variability of other hydraulic traits (Figure 6d), relaxing the constraints of maintaining a high *K_flower_* may have allowed morphological traits to vary more widely. Although this hypothesis has not yet been rigorously tested, some preliminary evidence suggests that morphological and physiological traits are correlated in flowers (Roddy, 2015).

### Implications for flower biomechanics

The variation in the hydraulic traits presented here have important implications for the structure and biomechanical performance of flowers. The low dry mass per area of flowers and their high *SWC* [Roddy et al. (2016); Figure 2] suggests that flowers may remain upright due to a hydrostatic skeleton maintained by turgor pressure rather than on a rigid, carbon-based skeleton. Relying on turgor pressure and a hydrostatic skeleton would increase the susceptibility of floral attraction to water limitation, which may be one explanation for why intraspecific variation in flower size is strongly influenced by water availability (Lambrecht & Dawson, 2007; Lambrecht, 2013). Although losing water is often considered expensive, for structures as ephemeral as flowers, the poor conversion rate of carbon for water (approximately 400:1; Nobel & others, 2005) may overwhelm the benefit of investing in long-lived carbon support structures, allowing flowers to be cheaper in terms of carbon though making them more vulnerable to drought-induced failure. Like in leaves with low mass per area (Scoffoni *et al*., 2014), the lower modulus of elasticity observed in flowers may predict the amount of tissue shrinkage as water potential declines and the failure of the hydrostatic skeleton. However, relying on turgor pressure to keep corollas upright is not the only method flowers may use to remain on display. Unlike leaves, floral corollas are often not planar, and many petals are curved or fused, which is a common way of increasing flexural stiffness independently of the modulus of elasticity (Vogel, 2013). Although the results presented here are only suggestive of the possible biomechanical strategies and tradeoffs flowers may use, linking the morphological, physiological, and biomechanical aspects of variation in floral form could yield novel insights into the multiple constraints acting on flowers and the evolution of floral innovations to overcome them.

## Acknowledgments

The authors thank K. Richardson and F. Rosin of the Arnold Arboretum and H. Forbes of the UC Botanical Garden for facilitating access to plant material. K. Niklas provided valuable input on interpretation. Jinyan Lei, SuYuan Li, and YiChan Li provided assistance with data collection. ABR was supported by a grant from the Yale Institute for Biospheric Studies.

